# The structure of Macrophage Expressed Gene-1, a phagolysosome immune effector that is activated upon acidification

**DOI:** 10.1101/580712

**Authors:** Siew Siew Pang, Charles Bayly-Jones, Mazdak Radjainia, Bradley A. Spicer, Ruby H.P. Law, Adrian W. Hodel, Sue M. Ekkel, Paul J. Conroy, Georg Ramm, Hariprasad Venugopal, Phillip I. Bird, Bart W. Hoogenboom, Ilia Voskoboinik, Yann Gambin, Emma Sierecki, Michelle A. Dunstone, James C. Whisstock

**Affiliations:** ARC Centre of Excellence in Advanced Molecular Imaging; Biomedicine Discovery Institute, Department of Biochemistry and Molecular Biology, Monash University, Melbourne Australia, 3800; FEI Company, Achtseweg Noord 5, Building 5651 GG Eindhoven, The Netherlands; London Centre for Nanotechnology, University College London, London WC1H 0AH, UK; Institute of Structural and Molecular Biology, University College London, London WC1E 6BT, UK; Department of Physics and Astronomy, University College London, London WC1E 6BT, UK; Cancer Immunology Program, Peter MacCallum Cancer Centre; Department of Genetics, The University of Melbourne, Parkville, Victoria 3010, Australia; School of Medical Science, Faculty of Medicine, University of New South Wales, Randwick; EMBL Australia, Single Molecule Node, Sydney Australia, 2052; EMBL Australia, Monash University, Melbourne Australia, 3800

**Keywords:** MPEG-1/Perforin-2, MACPF, Cryo-electron microscopy, Macrophage, Pore forming proteins, Immunity

## Abstract

Macrophage Expressed Gene-1 (MPEG-1; also termed Perforin-2) is an endosomal / phagolysosomal perforin-like protein that is conserved across the metazoan kingdom and that functions within the phagolysosome to damage engulfed microbes. Like the Membrane Attack Complex and perforin, MPEG-1 has been postulated to form pores in target membranes, however, its mode of action remains to be established. We used single particle cryo-Electron Microscopy to determine the 2.4 Å structure of a hexadecameric assembly of MPEG-1 that displays the expected features of a soluble pre-pore complex. We further discovered that the MPEG-1 pre-pore-like assemblies can be induced to perforate membranes through mild acidification, such as would occur within maturing phagolysosomes. We next solved the 3.6 Å cryo-EM structure of MPEG-1 in complex with liposomes. Remarkably these data revealed that a C-terminal Multi-vesicular body of 12 kDa (MVB12)-associated *β*-prism (MABP) domain interacts with target membranes in a mode that positions the pore forming machinery of MPEG-1 to point away from the bound membrane. This unexpected mechanism of membrane interaction raises the intriguing possibility that MPEG-1 may be able to remain bound to the phagolysosome membrane while simultaneously forming pores in engulfed bacterial targets.

---

Macrophage expressed gene-1 (MPEG-1, also termed Perforin-2) is one of the most ancient and highly conserved members of the Membrane Attack Complex / Perforin-like / Cholesterol Dependent Cytolysin (MACPF/CDC) superfamily, with orthologous sequences identifiable throughout the metazoa (1–4). Phylogenetic studies suggest that MPEG-1 represents the archetype of the human pore forming Membrane Attack Complex and perforin (5).

MPEG-1 is proposed to function within the macrophage phagolysosome(6–8). The Human protein comprises three regions; an ectodomain, a transmembrane domain that spans the vesicular membrane, and finally a short C-terminal cytosolic sequence (Supp. Fig. S1). The 636 amino acid ectodomain is contained within the vesicular lumen and contains an N-terminal MACPF/CDC domain together with a C-terminal region of unknown structure and function(4, 9, 10) (Supp. Fig. S1). While a bacteriocidal pore-forming function for MPEG-1 has not yet been definitively identified *in vivo* or *in vitro*, bacterial membranes derived from macrophages are decorated with ring-like structures(4). Further, some patients suffering from pulmonary nontuberculous mycobacterial infections carry MPEG-1 mutations(11).

Previous studies suggest that MPEG-1 ectodomain may be proteolytically cleaved away from the transmembrane domain(12). This region can also be expressed as a splice variant lacking the transmembrane / cytosolic region(8). We therefore expressed the human MPEG-1 ectodomain in order to investigate its function. Electron microscopy experiments revealed that the majority of recombinant material eluted as a stable multimeric complex (Fig. 1a). Accordingly, we used single particle cryo-EM to determine the structure of this assembly (Fig. 1b).

**Fig. 1.**
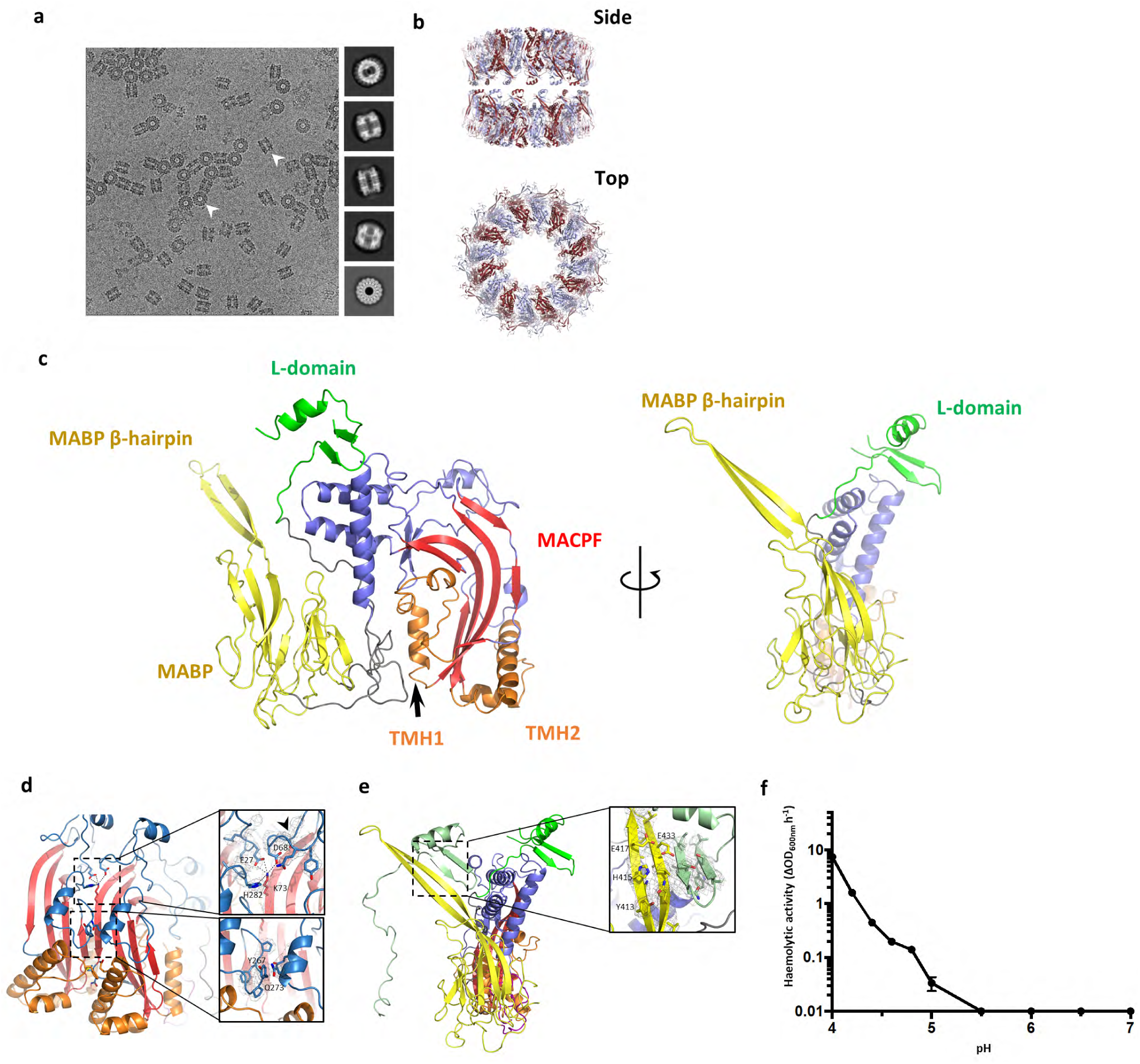
a) Cryo-EM image of MPEG-1 (top) together with a selection of class averages are shown. b) 2.4 Å structure of the MPEG-1 assembly; two hexadecameric rings (monomers alternately coloured) stack together to form a head to head dimer. c) Two views of the MPEG-1 monomer are shown (side on [left] and the C-terminal region (foreground) viewed from peripheral region of the assembly [right]). The MACPF/CDC domain is in blue, with the central sheet in red, and TMH-1/2 in orange. The central globular domain (MABP domain) is in yellow, and the L-domain is in green. d) Interactions made in the lumen, around the central *β*-sheet. e) Interactions made in trans by the *β*-hairpin (yellow) with the L-domain (green). f) Lytic activity of the pre-pore material at a range of different pHs (error bars represent SE, n = 4).

The 2.4 Å resolution structure of MPEG-1 reveals two hexadecameric rings stacked together in a head-to-head interaction (Fig. 1b). Three-dimensional classification experiments reveal that the rings are loosely associated with respect to each other (Supp. Fig. S2). Similar “double ring” structures have been observed in other pore forming proteins (e.g. aerolysin; gasdermin) and can form as a consequence of two membrane interacting surfaces interacting(13, 14).

The structure revealed that each MPEG-1 monomer within the 16-subunit assembly comprises an N-terminal MACPF/CDC domain, a central Multi-vesicular body of 12 kDa (MVB12)-associated *β*-prism (MABP) domain and a C-terminal region (termed the L-domain) (Fig. 1c). The MPEG-1 MABP domain possesses <5% primary amino acid sequence similarity to other homologues and was identified using Dali searches(15). The L-domain is largely extended, and includes a short 2 stranded *β*-sheet capped by an *α*-helix. Many MACPF/CDC proteins are produced as soluble monomers that bind to membranes via ancillary domains and then assemble into arc or ring-shaped pores (Supp. Fig. S3)(16–18). For many MACPF/CDCs, pore formation proceeds via assembly of a transient, intermediate pre-pore state, in which the membrane spanning regions have not yet been released(19–21). A conformational change within each MACPF/CDC domain in the pre-pore assembly then permits two small clusters of helices within each MACPF/CDC domain (termed transmembrane hairpin [TMH]-1 and -2) to unravel and penetrate the membrane as amphipathic *β*-hairpins (Supp. Fig. S3)(18, 22). The final pore comprises a large (compete or incomplete) *β*-barrel that spans the membrane(19, 23).

In contrast to other MACPF/CDCs studied to date(19, 24–27), the recombinant form of MPEG-1 has pre-assembled into a soluble oligomer that exhibits the expected features of a pre-pore assembly. Within each subunit, both TMH-1 and -2 are folded as small helical bundles (Fig. 1c). A central feature of the assembly is a 64-stranded *β*-barrel that represents the top, preformed portion of the future pore. The majority of MACPF/CDC domain mediated inter-subunit interactions are formed around the central lumen of the *β*-barrel (Fig. 1d, Supp. Table S1). Around the outside of the assembly both the MABP domain and the L-domain also mediate extensive inter-subunit interactions (Fig. 1e). The L-domain forms extensive inter-subunit interactions in trans with an extraordinary extended *β*-hairpin that extends from the MABP domain of the adjacent subunit (the MABP hairpin; Fig. 1c-e; Supp. Table S1). Finally, we note that one of the mutations associated with human disease (T73A(11); corresponding to T56 in our structure) maps to a loop buried in the inter-subunit interface.

We next investigated whether this MPEG-1 pre-pore could be induced to form pores. At neutral pH the recombinant material possessed no detectable activity in RBC lysis assays (Fig. 1f). However, we found that MPEG-1 started to gain membranolytic activity at acidic pH (*<*5.5) (Fig. 1f). To date, only one other acid activated MACPF/CDC protein has been identified - the bacterial cytolysin listerolysin O (LLO)(28, 29). The latter protein possesses enhanced pore forming activity at low pH and functions to facilitate pathogen survival via damaging the phagolysosome(30).

We reasoned that the MABP domain, a fold known to bind lipids, likely functions to localize MPEG-1 to target membranes. Previous studies reveal that MABP proteins interact with acidic membranes via a positively charged loop(31). Superposition of the MABP domain of MPEG-1 with the archetypal homologue from the Endosomal Sorting Complexes Required for Transport (ESCRT)-I MVB12 subunit reveal, however, that the membrane-binding loop maps to the extended MABP *β*-hairpin (Fig. 2a). These data thus suggested that the MABP domain is inverted with respect to the membrane binding domains of conventional MACPF/CDC proteins such as perforin and LLO (Supp. Fig. S3,S4).

**Fig. 2.**
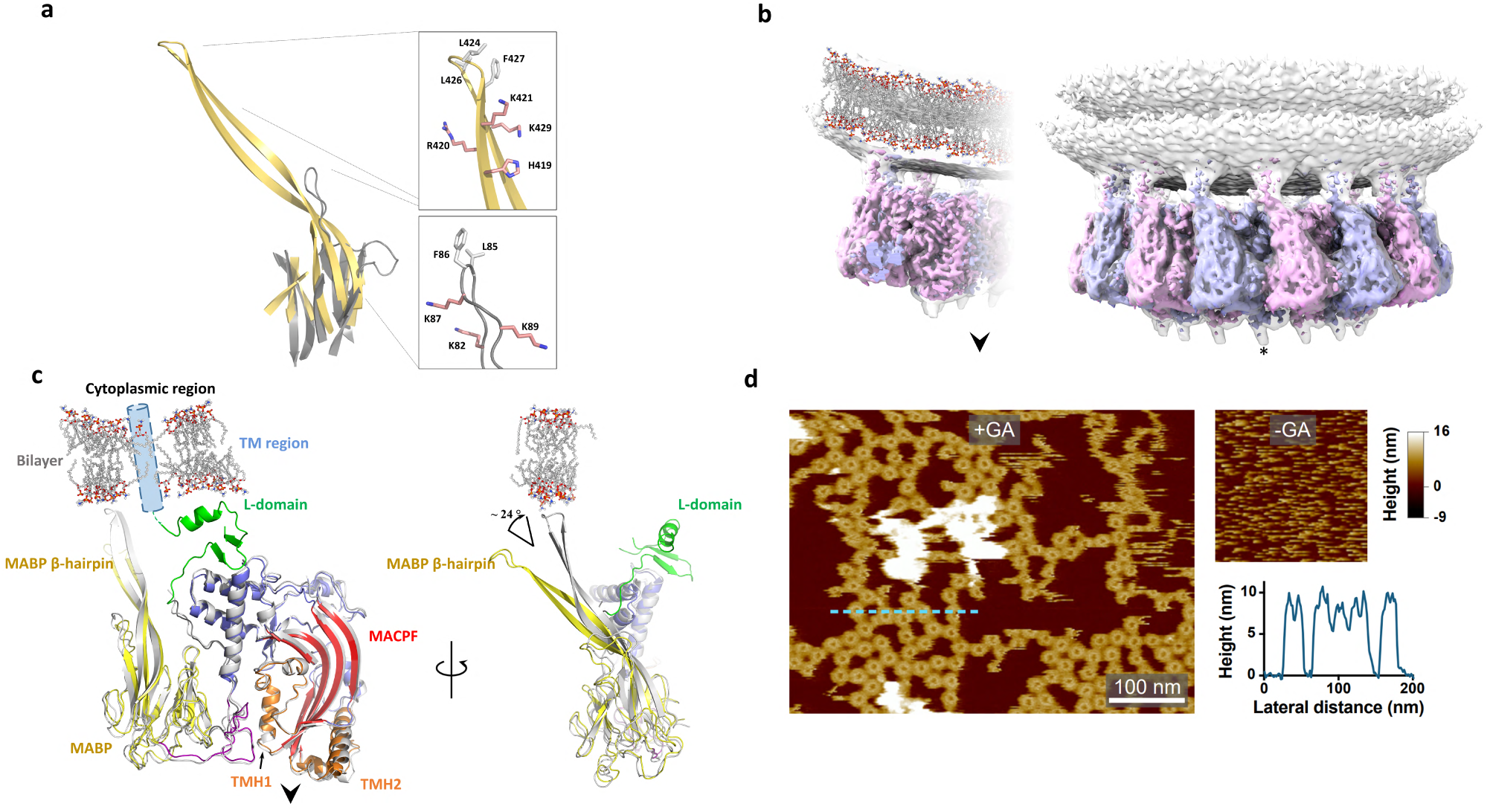
a) Structural superposition of the MPEG-1 MABP domain (yellow) and the MVB12-associated *β*-prism (grey). The lipid binding loop of the MVB12-associated *β*-prism maps to the MABP *β*-hairpin. The tip of both hairpins similarly contain a group of positively charged and hydrophobic residues. b) 3.6 Å structure of MPEG-1 bound to lipid membranes (subunits alternately coloured). A glycan moiety (indicated by an asterisk) is attached to TMH-2. The direction that the pore forming *β*-hairpins are released to form a membrane spanning *β*-barrel is shown by an arrow. c) Structural superposition of the MPEG-1 monomer derived from the head-to-head assembly (coloured as in 1c), and the lipid bound form (grey). The predicted position of the transmembrane domain (absent in the structure) is shown (blue cylinder). The *β*-hairpin shifts ∼ 24° in response to lipid interaction and the L-domain is disordered in the lipid bound form. The approximate position of the membrane is shown for reference. d) AFM images of MPEG on supported lipid bilayers consisting of *E. coli* lipid extract without (-GA) and with (+GA) glutaraldehyde fixation, at neutral pH. Scale bar: 100 nm. Colour scale: -9 to 16 nm.

Given these findings we sought to understand how MPEG-1 prepores coordinate membranes and determined the 3.6 Å cryo-EM structure of the assembly bound to liposomes at neutral pH (Fig. 2b). These data reveal that the MABP domain binds lipids in a canonical fashion through the MABP hairpin. The latter region bends as a consequence of interactions with the membrane, a shift that involves breaking contacts with the L-domain. Indeed, the C-terminal portion of the L-domain is not visible in electron density, suggesting that this region becomes flexible when the MABP hairpin distorts through membrane interactions. (Fig. 2c). The remainder of the structure appears essentially unaltered (Fig. 2b-c). Atomic force microscopy (AFM) studies also confirms the binding of MPEG-1 prepores to membrane (Fig. 2d). Consistent with previous studies on perforin (and CDC) prepores on supported lipid bilayers(32), MPEG-1 prepores appear as mobile (and therefore poorly resolved) features on the membrane, which upon fixation with glutaraldehyde can be clearly identified as ring-shaped assembles of dimensions consistent with the EM data (Fig. 2d).

The observed mode of MPEG-1 membrane binding is also consistent with the location of the C-terminal transmembrane region (absent in our construct; Fig. 2c), which in full-length protein immediately follows the L-domain. Collectively, these data suggest that MPEG-1 includes two independent features - a transmembrane sequence and the MABP hairpin - that could each function redundantly to ensure localisation of the protein to the inner leaf of the vesicular membrane. Such an interaction mode, if it were to be maintained during lytic function, would mean that the pore forming regions released by the MACPF domain would point directly away from membrane coordinated by the MABP domain (Fig. 2b, c). The implication of our findings is therefore that MPEG-1 may be able to bind one membrane system through the MABP domain while simultaneously forming pores in a second membrane system. In support of this idea, analysis of the liposome dataset collected at neutral pH revealed occasional examples of MPEG-1 apparently coordinated to one liposome, while forming pores in a second (Fig. 3). However, there were only a few examples of such complexes present (typically 1-2 pores / image) and it was not possible to collect a dataset. Further, despite extensive efforts, incubation of MPEG-1 / liposome mixtures at acidic pH, failed to yield a sample suitable for collection of a high-quality dataset, with pore-containing liposomes typically clumped together.

**Fig. 3.**
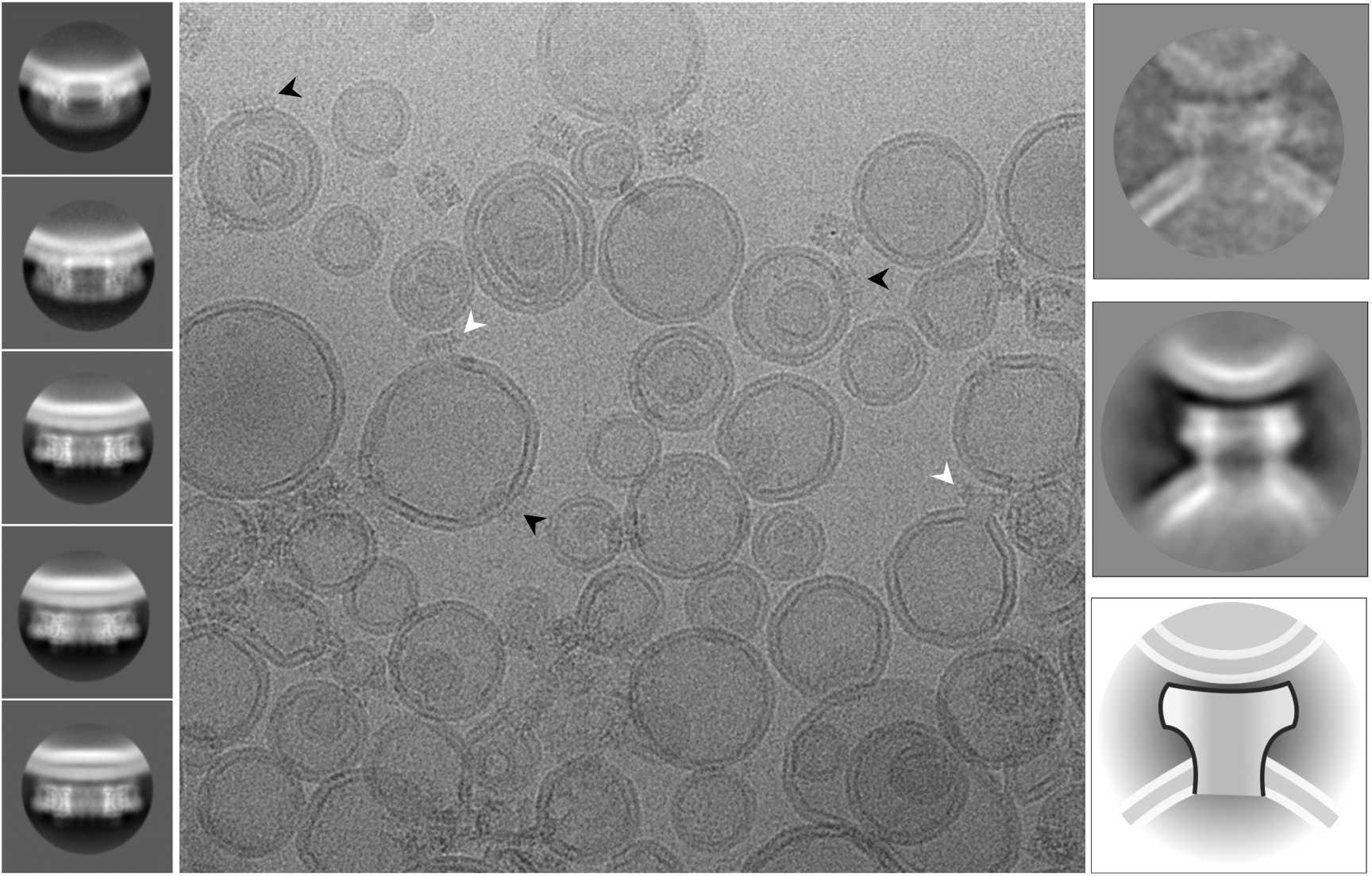
Examples of MPEG-1 bound to liposomes reveal single rings bound in the pre-pore state (black arrows) and the occasional example of pores (i.e. the membrane is absent in the pore lumen), that coordinate a second liposome through the top portion of the ring (white arrow). We were not able to identify examples of pores bound to a single liposome, i.e. all pore structures we observed bridged between two liposomes. Class averages for pre-pores (left) and pores (right) are shown.

In conclusion, our data lends support to the idea that MPEG-1 is an acid activated protein that functions to form pores in bacteria located within the phagolysosomal compartment(7). Our data further reveals the intriguing finding that at neutral pH the MABP domain binds membranes in an orientation that would preclude MACPF pore formation in the MABP-bound membrane (Fig. 2b, c). This contrasts other MACPF proteins such as perforin and the CDCs, which deploy ancillary lipid binding domains in order to assemble on the membrane targeted for pore formation (Supp. Fig. S3).

Given these findings, we speculated how MPEG function might form pores in engulfed targets. One possibility is that other intra-cellular events, for example proteolysis(12) and / or changes in pH (reported here), permits release of MPEG-1 into the lumen of the phagosome where it may function to form pores in a conventional perforin / CDC-like fashion (Fig. 4a). Alternatively, we suggest that the observed mode of MABP membrane binding may permit interaction with the phagolysosome membrane while simultaneously forming pores in engulfed bacteria (Fig. 4b). Indeed, such a mechanism may be advantageous in regards to preventing unwanted auto-lysis of the phagolysosome.

**Fig. 4.**
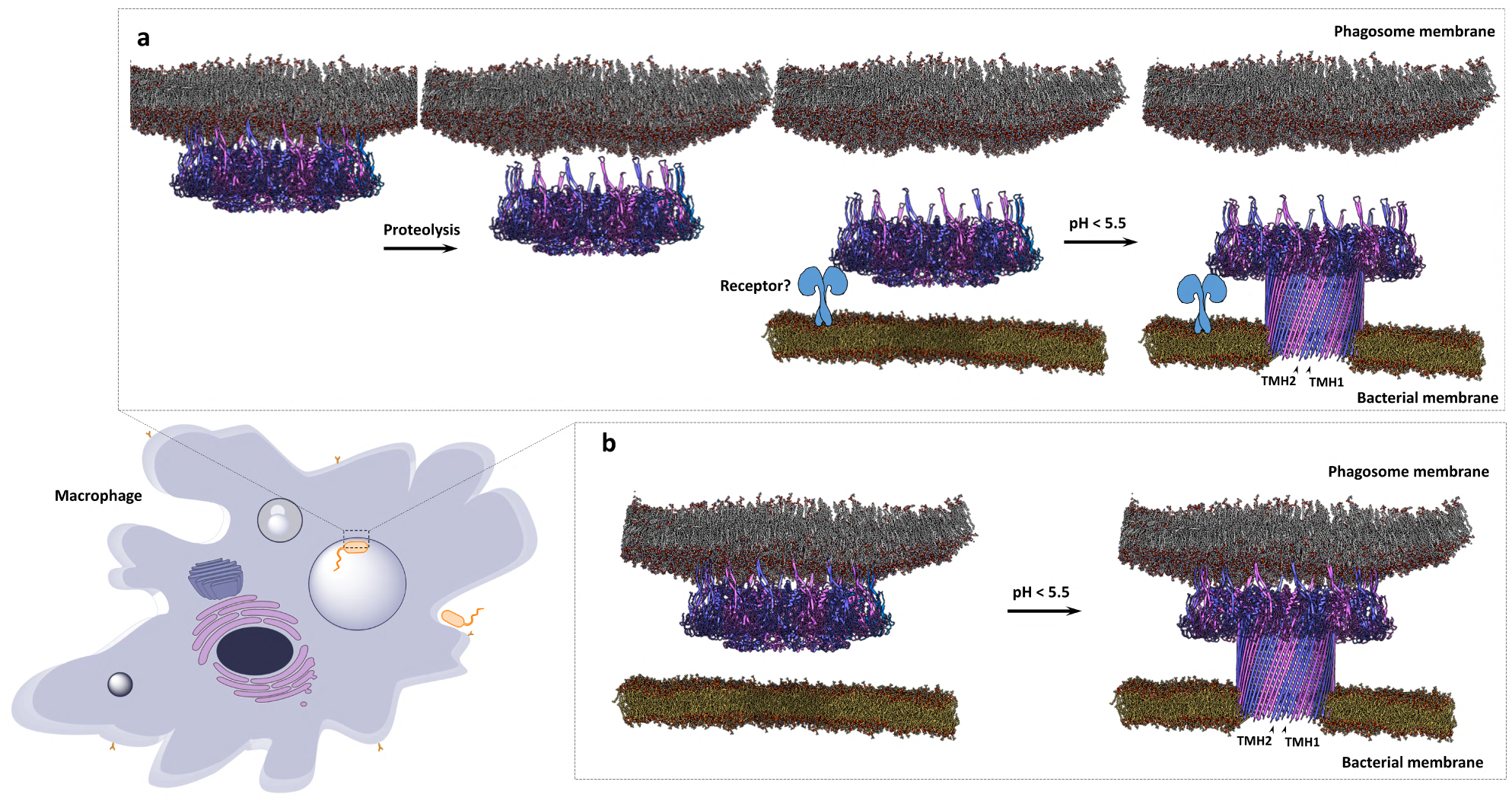
Schematic illustrating two proposed mechanisms for control of MPEG-1 function. a) MPEG-1 prepores (alternative subunits coloured) are localized to the phagolysosome membrane, either through the transmembrane helix, or the MABP *β*-hairpin. Upon acidification or proteolytic activity, the pre-pores are released from the membrane into the phagosome lumen. Via an unknown means these recognise engulfed target membranes and form pores. b) Alternatively, prepores remain tethered to the phagosome membrane inner leaflet via the MABP *β*-hairpin and are triggered to form pores in engulfed bacteria upon acidification.

## Methods

### Protein purification

The human MPEG-1 gene for the soluble ectodomain (NCBI-gene ID: 219972, nucleotide 49-1959) was synthesised (Genescript) with codon usage optimised for insect cell expression. The native signal peptide was replaced with the honeybee melittin signal peptide for protein secretion, and a C-terminal hexahistidine tag was introduced for affinity purification. The recombinant gene was cloned into pFastBac1 expression vector (Life Technologies) for bacmid production. Baculoviral stocks (P1 to P3) were generated in Sf9 or Sf21 insect cells as described in the supplier’s protocols.

For MPEG-1 expression, Sf21 cells (2 × 10^6^ cells mL^−1^) were infected with the P3 viral stock in Insect-XPRESS protein-free medium (Lonza). The infected insect cells were grown at 27 °C with shaking for 66 hours. The insect cell supernatant was harvested by centrifugation at 1000 × g for 10 minutes. The clarified supernatant was cooled to 4 °C and buffer exchanged into 20 mM Tris-HCl, pH 8.0, 300 mM NaCl, 20 mM imidazole by extensive dialysis or by tangential flow filtration (Cogent M1 TFF system, Millipore).

The buffer-exchanged supernatant was clarified by filtrating through a 0.8 µm membrane before loading onto a Ni-NTA agarose column (Qiagen). The column was washed with buffer containing 40 mM imidazole and the His-tagged MPEG-l eluted with buffer containing 500 mM imidazole. The protein peak fractions were pooled and dialysed against 20 mM Tri-HCl, pH 7.2, 300 mM NaCl, 10 % (w/v) glycerol at 4 °C overnight. The pooled protein was further purified by size-exclusion chromatography on a Superose 6 10/300 column (GE Healthcare Life Sciences). MPEG-1 fractions were monitored using SDS-PAGE, haemolytic activity assay and negative-stained EM. For further cryoEM work, MPEG-1 fractions were selected and pooled carefully based on negative-stained EM analysis for sample homogeneity. For all other work, fractions are assessed and pooled based on purity on SDS-PAGE and haemolytic activity. Pooled fractions were concentrated to 0.5 to 1.0 mg mL^−1^ and snap-frozen in liquid nitrogen for storage at - 80 °C.

A similar approach was used to produce a mutant form of MPEG-1_*L*425*K*_. While the structures yielded by this mutant are not discussed extensively in this paper, the information derived from this structure proved important for determining the complete structure of wild type protein.

### Red blood cell (RBC) lysis assay

Rabbit blood was obtained from Applied Biological Products Management and collected in the presence of heparin to prevent clotting. To prepare the RBCs for the lysis assays, the blood was firstly fractionated by centrifugation at 3000 × g at 4 °C for 15 min to pellet the RBC. The cells were resuspended gently and washed three times with equal volume of HBS. The final RBC pellet was resuspended in preservative Celpresol (CSL™) to the original blood volume for storage at 4 °C. Before each lysis assay, the rabbit RBC were pelleted and washed in wash buffer (5 mM HEPES pH 7.0, 75 mM NaCl, 2.5 % (w/v) glucose, 0.15 mM CaCl2, 0.5 mM MgCl_2_) three times to remove the preservative solution, any lysed cells and soluble haemoglobin. MPEG-1 membranolytic activity was monitored by reduction of turbidity of RBC suspension at OD600nm. The washed red blood cell was diluted 20-fold into the reaction buffers (20 mM buffer, 75 mM NaCl, 2.5 % (w/v) glucose) with the respective pH (sodium acetate buffer for pH 4.5-5.5, MES buffer for pH 6.0-6.5 and Tris-HCl for pH 7.0) to give a starting OD600nm 1.5-2.0. The haemolytic activity was initiated by adding the diluted RBC to MPEG-1. MPEG-1 membranolytic activity is calculated by the rate of RBC lysis (ΔOD600nm h^−1^) per *µ*g of MPEG-1 protein. All reactions were carried out in duplicate and blanked against identical assays except without the addition of MPEG-1, four independent experiments were performed (n = 4).

### CryoEM sample preparation and data collection

For cryo-EM of MPEG-1, the protein sample was buffer-exchanged into 20 mM Tris-HCl, pH 7.2, 300 mM NaCl to remove the glycerol, and concentrated between 2.0 to 2.5 mg mL^−1^. The cryo-EM girds were generated using the Vitrobot System (Thermo Fischer Scientific). Initial grid freezing conditions were tested and screened on a Tecnai T12 electron microscope (Thermo Fischer Scientific). Briefly, 3 µL of MPEG-1 were applied to a glow-discharged QUANTIFOIL Cu R 1.2/1.3 grid, with blotting conditions as followed. The temperature was set to 4 °C with the relative humidity option turned off, the grids were blotted with a blot time of 2.5 s, blot force of −1 and drain time of 1 s. The grids were snap-frozen in liquid ethane and stored under liquid nitrogen until TEM data collection. For cryo-EM of liposome / MPEG-1 complex, the protein solution was prepared as described above into HEPES buffered saline (HBS, 20 mM HEPES pH 7.0, 150 mM NaCl). The liposomes were composed of POPC and POPS (Avanti Polar Lipids) in equal ratio. To prepare the liposomes, 2 mg of each POPC and POPS in chloroform stock solutions were mixed and dried under argon in a clean test tube, and desiccated under vacuum for at least 4 hours. The dried lipid mixture was resuspended in 0.5 mL HBS buffer by vortexing for 1 minute, then snap-frozen in liquid nitrogen, and sonicated in a warmed (30 °C) ultrasonic bath for 15 minutes. This process was repeated three times. The lipid suspension was then extruded using a polycarbonate membrane with pore size of 0.1 *µ*m (Avanti Polar Lipid) to obtain unilamellar liposomes. To generate liposome / MPEG-1 complex, 10 *µ*L POPC:POPS liposomes was mixed with 5 *µ*L MPEG-1 (1.7 mg mL^−1^) and incubated at 37 °C for three hours. The liposome and MPEG-1 mixture was frozen onto a glow-discharged QUANTIFOIL Cu R 2/2 grid as described above with the following modifications. The Vitrobot conditions were set to 22 °C with 100 % humidity, the grids were blotted with a blot time of 2.5 s, blot force of − 5 and drain time of 1 s.

Data collection parameters have been summarised (Supp. Table S2) for each data set. Briefly, dose fractionated movies were collected on a Titan Krios (Thermo Fischer scientific), equipped with a Gatan Quantum energy filter (Gatan) and either a Falcon II (Thermo Fischer scientific) or Summit K2 (Gatan) direct electron detector. Data acquisition was performed using either SerialEM(33) or EPU (Thermo Fischer Scientific).

### CryoEM image processing

Upon finalisation of data collection to calibrate pixel size and estimate magnification anisotropy, images of gold diffraction grating (Agar scientific) were collected at the same magnification as the collection. Analysis performed with mag_distortion_estimate(34) indicated magnification anisotropy was indeed present (Supp. Table S2). Therefore, dose fractionated movies were corrected for beam induced motion, anisotropy and radiation damage within MotionCor2(35). Super resolution movies were additionally down sampled by a factor of 2, applied by Fourier cropping within MotionCor2. All aligned movie frames were subsequently averaged into dose-weighted and non-weighted sums for further processing.

Particle coordinates were determined using various software depending on the data set, a combination of gAutomatch (Zhang et al, unpublished; https://www.mrc-lmb.cam.ac.uk/kzhang/Gautomatch/), crYOLO (v1.1) (Wagner et al., 2018. bioR*χ*iv doi:10.1101/356584) and manual picking were employed. A rapid single round of 2D classification in cryoSPARC(36) (v1) was employed to remove contaminants and poorly picked areas. CTF estimation of whole non-dose weighted micrographs was initially performed with CTFFIND4 (v4.1.10)(37). On-the-fly processing was performed in RELION (v2.1)(38) to assess data quality. Initial models were all generated ab initio in either cryoSPARC or in EMAN(39) (v2.2) by the common line method for the MPEG-1_*W T*_ dataset (Supp. Fig S2).

All further processing was performed in RELION, unless otherwise stated. Particles belonging to clean classes were then subjected to 3D classification to remove malformed particles and projections with poor signal-to-noise. Clearly abnormal classes were discarded and 3D refinement was per-formed on the remainder. Per-particle CTF parameters were refined without further alignments in cisTEM (v1.0b)(40) until convergence, here the refinement resolution was maintained above the fall off of the FSC. Particles with poor scores from cisTEM were discarded.

Masked 3D classification with the previously refined angular and CTF parameters without additional alignments was carried out for all datasets to identify homogenous subpopulations. In the case of the soluble MPEG-1_*L*425*K*_data set, this resulted in two distinct populations. One population exhibiting a discrete D16 conformational state, termed the *β*-conformation, consisted of approximately one sixth of all the particles (Supp. Fig S8). This conformer displayed isotropic resolution and enabled the MABP domain to be resolved. The second population, termed the *α*-conformation, exhibited continuous conformational heterogeneity. This population could be further reduced into a continuous distribution of D16 states. Here conventional refinement failed, resulting in reconstructions that had radially decreasing quality where only the central most region was resolved. Analysis by RELION (v3.0b2) multi-body procedure(41) resulted in a marked improvement in resolution. Indeed, principal component analysis highlighted additional degrees of freedom and loose association of each ring (Supp. Fig S2). Symmetry breaking hence occurred as a result of relative motion between each ring.

To overcome symmetry breaking, localised reconstructions were performed on each ring and additionally on individual monomers thereby reducing the symmetry to C16 and C1 respectively. Firstly, partial signal subtraction of both the top and bottom rings was performed in RELION to obtain two sub-particles per image. These were individually refined in cryoSPARC with dynamic masking. Localised reconstruction resulted in a notable improvement of FSC and quality of the reconstruction. Later sub-particles were merged into a single data set, which refined to the highest resolution in the central region of the complex (MACPF domain). However, the quality of the reconstruction in the MABP domain remained uninterpretable due to intramolecular, conformational heterogeneity between subunits (Supp. Fig. S2,S7). Therefore, sub-particles corresponding to individual monomers were re-extracted along with partial signal subtraction using the localised reconstruction scripts(42). These sub-particles were aligned to a common axis and sorted by masked 3D classification without further alignments, where a subpopulation with substantially higher homogeneity was identified thereby enabling the full structured to be resolved (Supp. Fig. S2,S7).

In the case of the liposome / MPEG1 complex, density subtraction of the lipid bilayer was crucial for accurate alignments. A mask of the lipid bilayer was created by segmenting the best reconstruction with Segger (v1.9.5)(43). Density subtraction of the lipid bilayer was then performed in RELION followed by masked refinement(44). This was repeated for a total of two subtractions as residual membrane density was observed. The final map was reconstructed from the original particles with the optimised alignments.

Per frame B-factor weighting was performed by movie refinement and particle polishing of the final subset of particles after classification and refinement of all reconstructions, except in the cases of localised reconstruction where polishing was performed prior to sub-particle extraction. Lastly polished particles were re-refined. For reconstructions below 3 Å, additional corrections and refinements were performed as follow. Ewald sphere correction, astigmatism and beam tilt refinement were performed in in RELION (v3.0b2)(45). In the case of super-resolution images, the data were resampled to the original pixel size and the final iteration of refinement was continued in RELION. While these manipulations did not yield improvements for most reconstructions, Ewald sphere effects did appear to affect the MPEG-1_*L*425*K*_*α*C16 reconstruction, marginally improving the resolution by 0.04 Å.

Global resolution was calculated by the gold standard Fourier shell correlation (FSC) at the 0.143 criterion. Local resolution was estimated for all reconstructions in RELION using a windowed FSC_0.143_. For b-factor sharpening, MonoRes and LocalDeblur were used to re-estimate local resolution and enhance high resolution features by local sharpening respectively (Ramírez-Aportela et al., 2018. bioR*χ*iv doi:10.1101/433284). Locally sharpened maps were subsequently filtered by local-resolution with blocfilt(46) or with RELION. Analysis of resolution anisotropy was performed by 3DFSC(47). Any conversions between software were performed with EMAN (v2.2), code written in-house or by D. Asarnow and J. Rubeinstein.

### Model building and analysis

Phenix real-space refinement was performed on all models, followed by manual verification of Ramachandran values and fit-to-density within Coot(48). Analysis of model-to-map quality was performed by a combination of EMRinger(49) and MolProbity(50) scores. Lastly the map-to-model FSC was calculated in Phenix. All figures and visualisation of models, maps and trajectories were performed in UCFS ChimeraX(51), Pymol(52) or VMD(53). Analysis of contacts was performed by CONTACT in Coot and PISA(54).

Model building of the MACPF region was originally performed *de novo* into the MPEG-1_*W T*_ reconstruction in Coot, while the MABP region was poorly resolved and could not be built due to symmetry breaking. Later the MABP domain was clearly resolved in both the MPEG-1_*L*425*K*_*β* D16 and MPEG-1_*L*425*K*_*α* C1 reconstructions (Supp. Fig. S2). These maps were then used to build the remainder of the structure *de novo* in Coot. The MPEG-1_*L*425*K*_*α* C1 model was used as a template to make the final model of MPEG-1_*W T*_, this was fitted into the MPEG-1_*W T*_ reconstruction by rigid body docking followed by Phenix real-space refinement.

A model of the lipid bound MPEG-1 prepore was obtained by molecular dynamics flexible fitting. Here the reconstruction of liposome/MPEG-1_*L*425*K*_ was used as a biasing potential map and a single ring of the MPEG-1_*L*425*K*_*α* conformer was flexibly guided into the map by namd2/MDFF(55). As the L-domain becomes disordered upon lipid binding, this region was removed from the model.

### Atomic Force Microscopy

AFM experiments were using parameters, setups, and analysis as previously described (32). *E. coli* total lipid extract was purchased from Avanti Polar Lipids, Inc. From the lipid extract, small unilamellar vesicles with a nominal diameter of 50 nm were produced. The small unilamellar vesicles were obtained at a concentration of 1 mg mL^−1^ in 150 mM NaCl, 20 mM HEPES, pH 7.4. 6 µL of the small unilamellar vesicle solution were injected onto a freshly cleaved mica surface (⊘9.9 mm, Agar Scientific) covered in 80 µL of adsorption buffer (150 mM NaCl, 20 mM HEPES, 25 mM MgCl_2_, pH 7.4), and subsequently incubated for 30 min at room temperature to form an extended supported lipid bilayer film. Prior to injecting protein, the supported lipid bilayer was gently washed 15 times with 80 µL of the adsorption buffer to remove residual lipid vesicles. MPEG-1 was injected onto the lipid bilayer to a concentration of 70 nM and incubated for 15 min at 37 °C. The resulting membrane bound, mobile MPEG-1 assemblies were imaged with AFM. To immobilise membrane bound MPEG-1, the assemblies were cross-linked by addition of 0.12 % glutaraldehyde (TAAB Laboratories) and incubation for 10 min at room temperature, before imaging with AFM.

### Data Availability

Cryo-EM maps and atomic models will be deposited in the Electron Microscopy Data Bank (EMDB) under accession codes EMBD XXX, EMBD XXX, etc. Each EMDB entry includes five maps: (1) the low-pass-filtered map without amplitude correction as a default; (2) the low-pass-filtered map with amplitude correction by a negative B-factor shown in Extended Data Table 1; (3) the half maps without any post-processing such as low-pass-filtering and amplitude correction; and (4) the mask associated with refinement and sharpening. Coordinates are available from the RCSB Protein Data Bank under accession codes PDB XXX, PDB XXX, etc. Raw data are available from the corresponding author upon request.

## ACKNOWLEDGEMENTS

JCW is an Australian Research Council (ARC) Laureate Fellow and an Honorary National Health and Medical Research Council (NHMRC) Senior Principal Research Fellow. He acknowledges the previous support of an Australian Research Council (ARC) Federation Fellowship. MAD acknowledges the support of an ARC Future Fellowship and further acknowledges the previous support of an NHMRC Career Development Award. CBJ acknowledges the support of the Australian Government RTP scholarship. AWH acknowledges support by the UCL Grand Challenge scheme and the Sackler Foundation. BWH acknowledges support from the UK Biotechnology and Biological Sciences, and Engineering and Physical Sciences Research Councils (BBSRC, BB/N015487/1, and EPSRC, EP/M028100/1). We thank the staff of the Monash Ramaciotti Centre for Electron Microscopy, the Monash protein production and proteomics platforms, and the support of the MASSIVE supercomputer team. We thank Prof Helen Saibil and Natalya Lukoyanova (Birkbeck College, London) for helpful discussions. This preprint was formatted in LATEXusing an adaptation of Ricardo Henriques’ template.

---

**Fig. S1.**
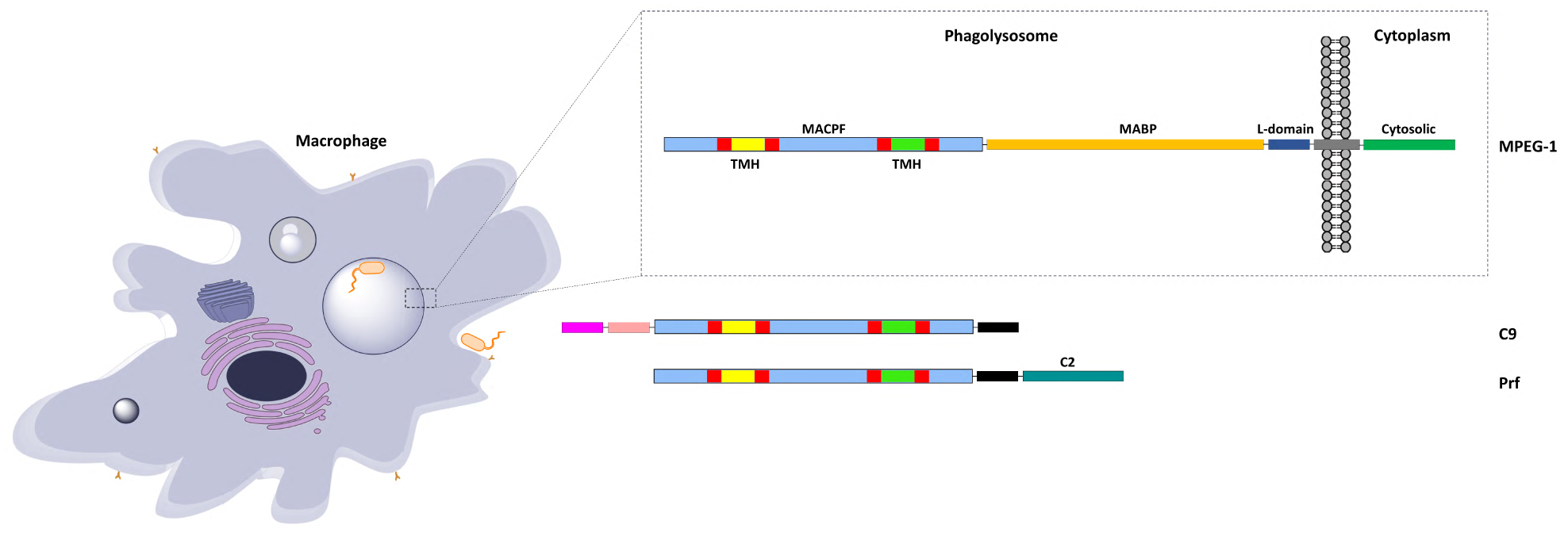
Schematic of MPEG-1 domain architecture and subcellular localisation. MPEG-1, complement C9 and perforin share a homologous MACPF domain, however only MPEG-1 possess a transmembrane tether and cytosolic region. The proposed orientation of MPEG-1 in the vesicular compartment places the pore forming domain in the acidic lumen of the phagosome, while the cytosolic region is exposed in the cytoplasm(4).

**Fig. S2.**
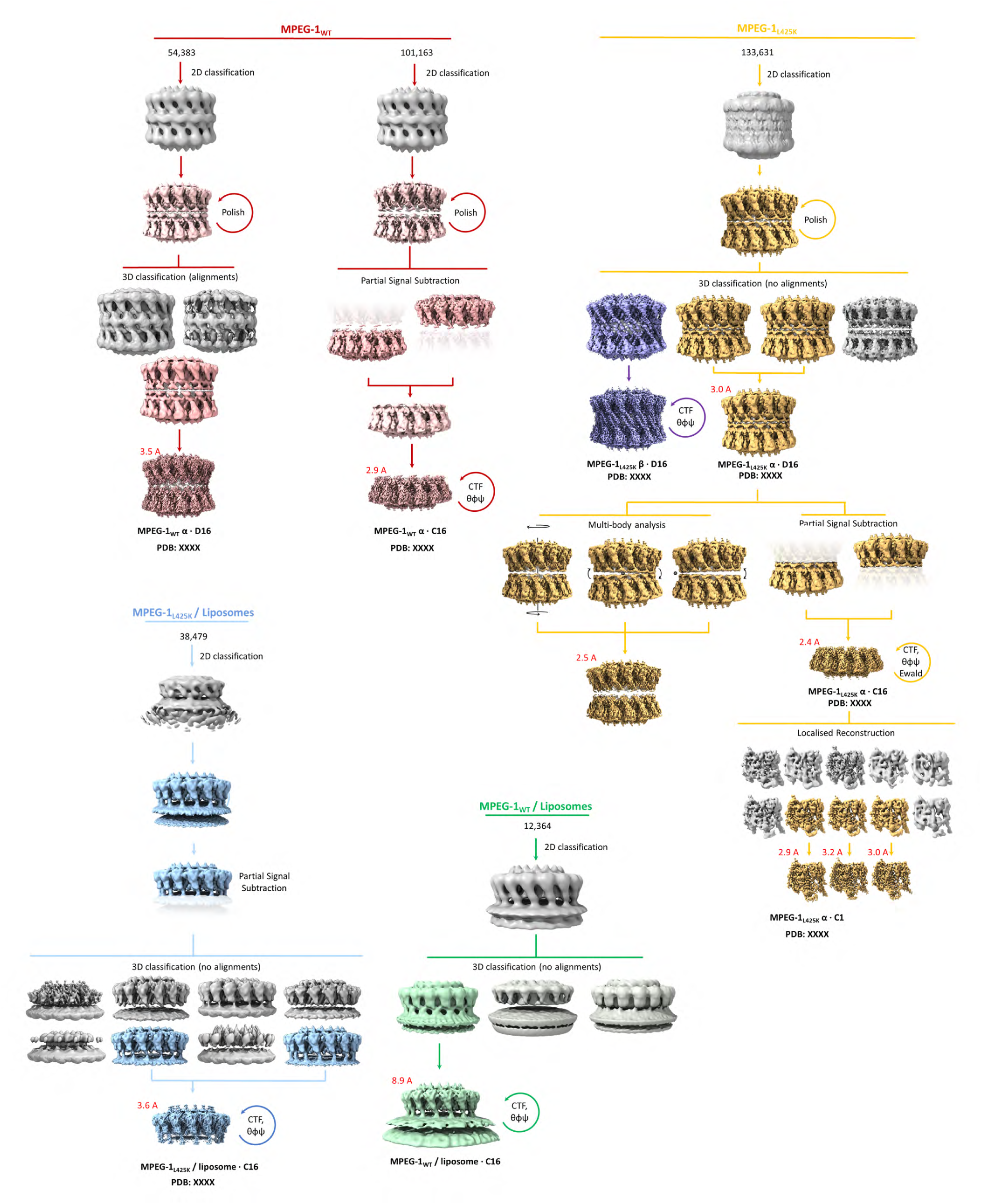
Summarised overview of the analysis and workflow for all four cryo-EM data sets. Two data sets of the soluble head-to-head assembly were collected, namely MPEG-1_*W T*_ (rose) and MPEG-1_*L*425*K*_ (yellow/purple). Multiple rounds of classification and multibody analysis of the MPEG-1_*L*425*K*_ dataset revealed discrete conformational heterogeneity, in addition to continuous conformation heterogeneity between the two ring systems. Furthermore, intra-molecular discrete conformational heterogeneity was present at the subunit level in the MABP domain necessitating localised reconstruction. Additionally, two data sets of the lipid bound pre-pore assembly were obtained, referred to as MPEG-1_*W T*_ / liposome (green) and MPEG-1_*L*425*K*_ / liposome (blue).

**Fig. S3.**
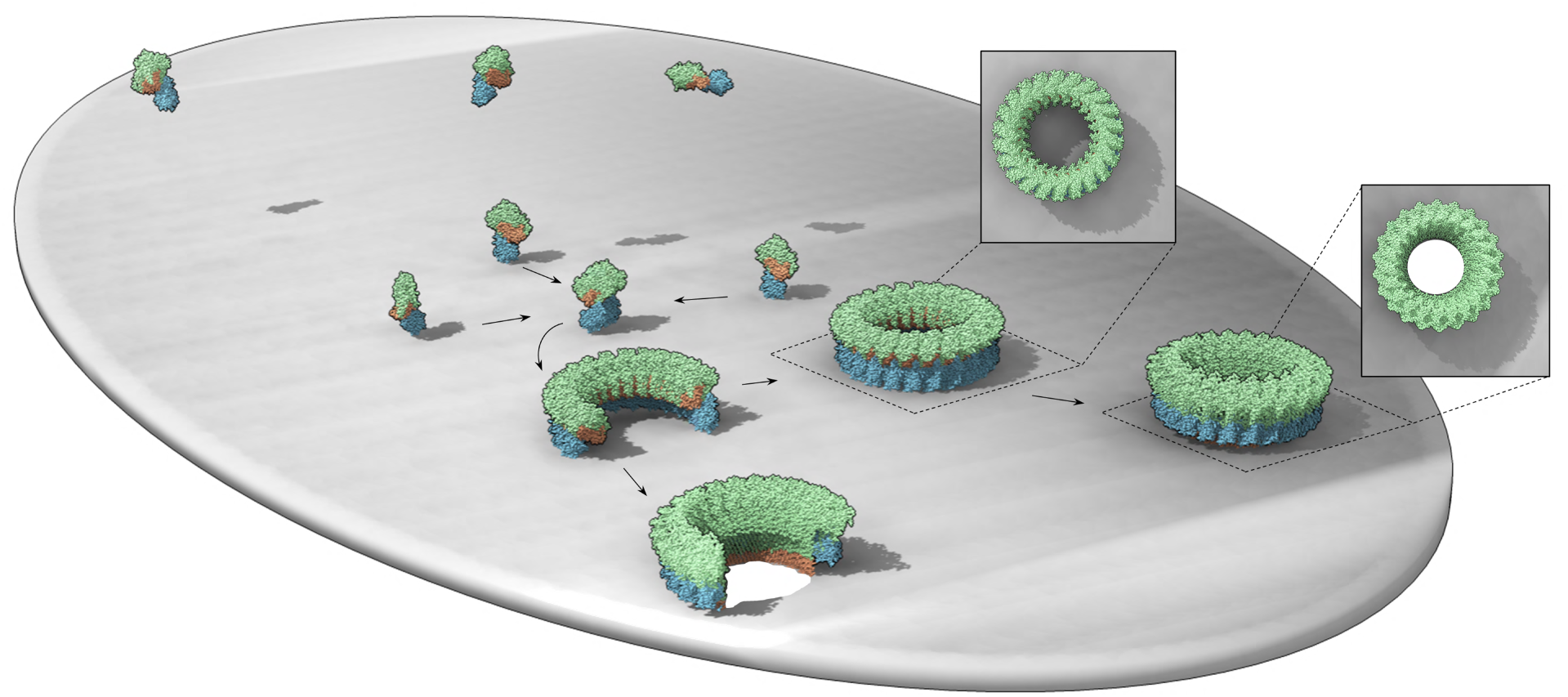
Simple illustration of the canonical MACPF and CDC mechanism of pore formation. Initially freely diffusing soluble subunits recognise and bind to target membranes by employing a target recognition domain (e.g. C2 in Perforin). Membrane bound monomers undergo two-dimensional diffusion along the surface of the membrane assembling into larger oligomers that have yet to insert into the membrane i.e. prepores. These prepores then undergo a conformation change whereby the TMH regions unfurl and insert into the membrane forming a giant (complete or incomplete) *β*-barrel pore. Insets represents a view from above.

**Fig. S4.**
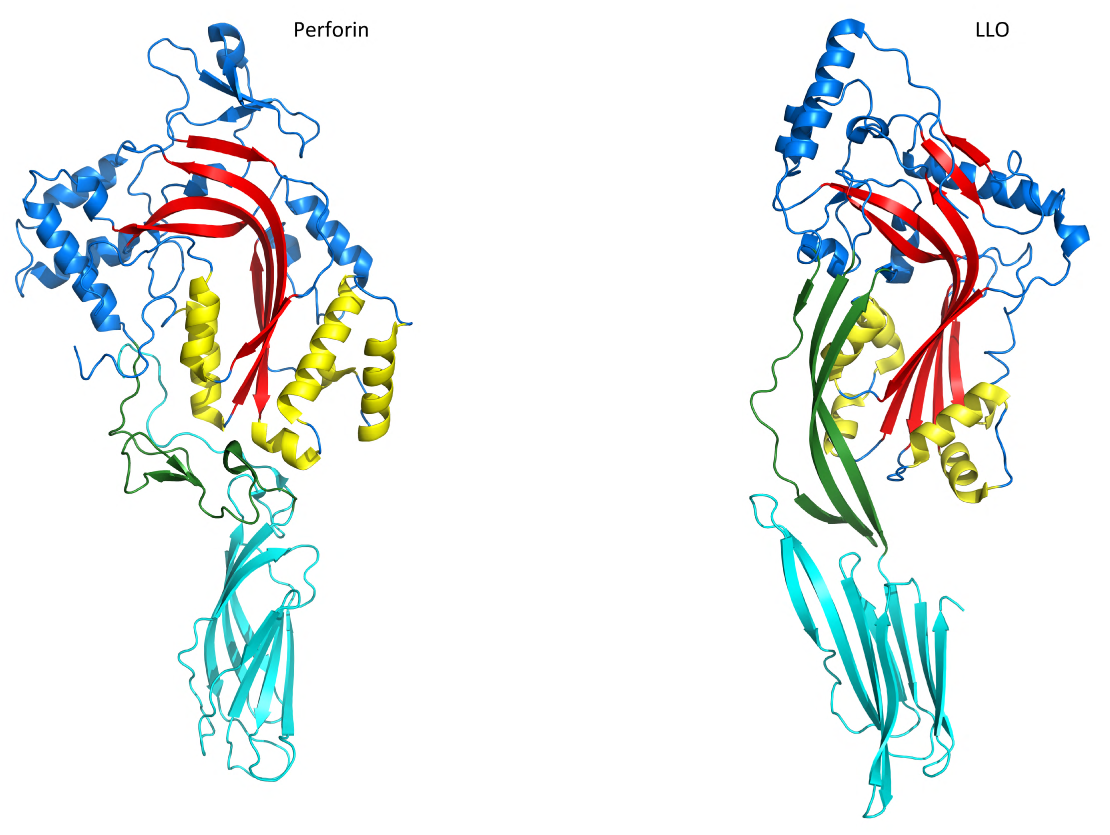
Structural comparison between lymphocyte Perforin and lysteriolysin O (LLO) highlighting the ancillary membrane binding domains (Perforin C2 domain and LLO domain 4; cyan). The pore forming MACPF/CDC domain is coloured blue, with the central b-sheet in red and TMH1/2 in yellow. Notably in these exemplars, the MACPF/CDC domain is oriented in the same direction as the ancillary targeting domain. The shelf domain, which in CDCs facilitates the collapse, is coloured green.

**Fig. S5.**
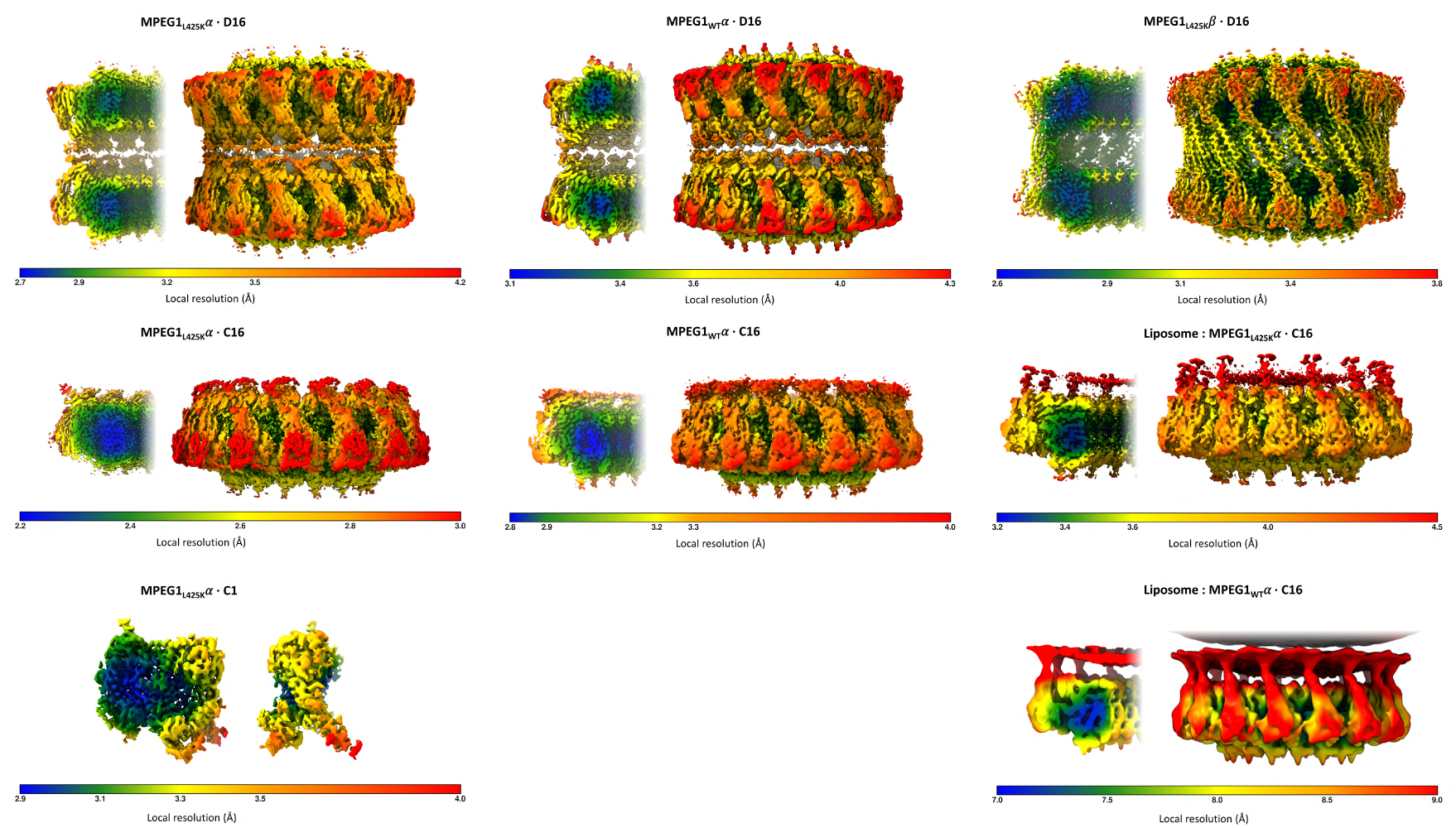
Local variation in resolution by gold-standard Fourier shell correlation (FSC) at 0.143. Local resolution ranges in Ångström (Å) are displayed as a colour gradient (blue to red) on the locally filtered, sharpened maps. Scales are provided below each reconstruction.

**Fig. S6.**
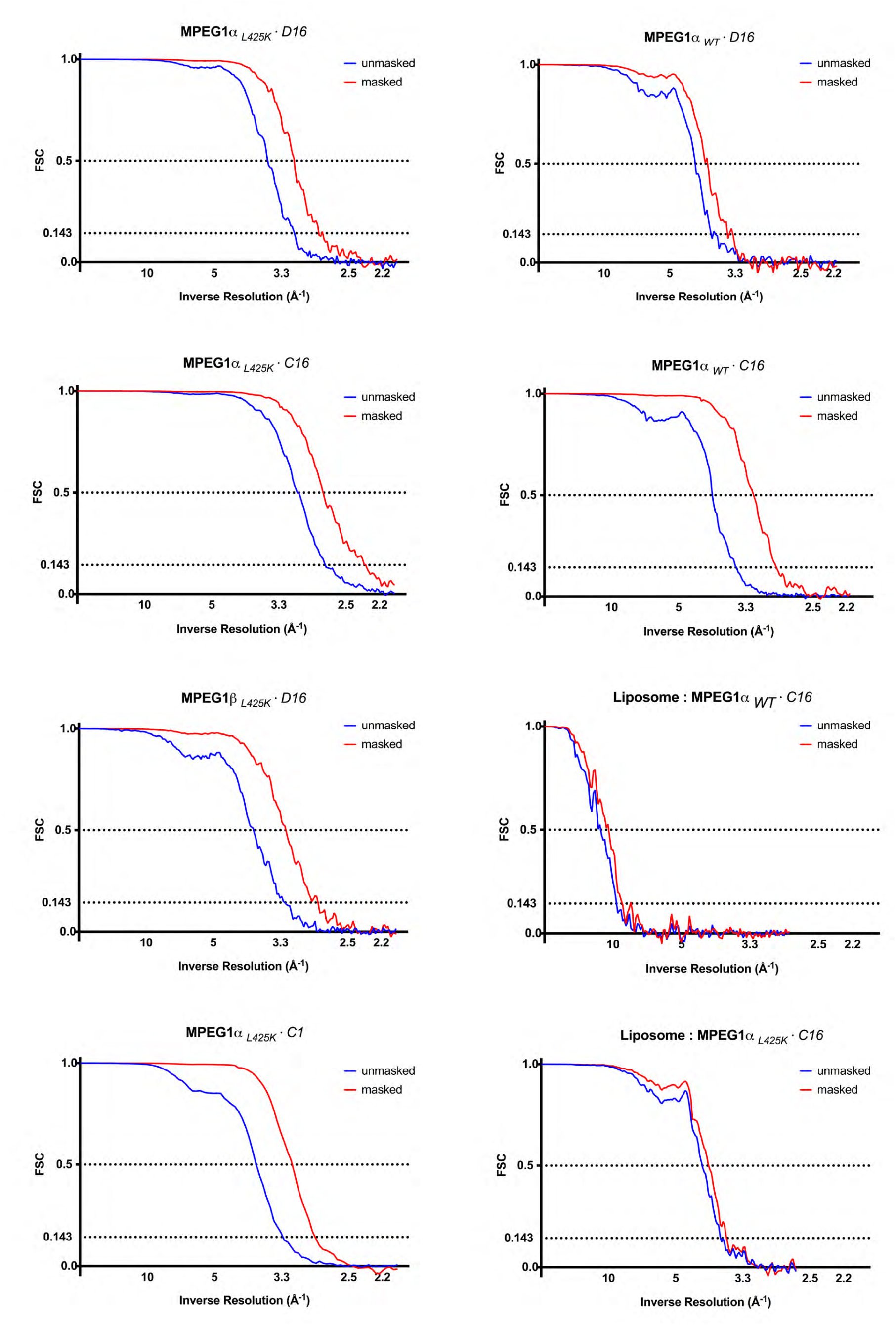
Global gold-standard Fourier shell correlation (FSC) of masked and unmasked maps. Two FSC criteria are plotted at 0.5 and 0.143 thresholds.

**Fig. S7.**
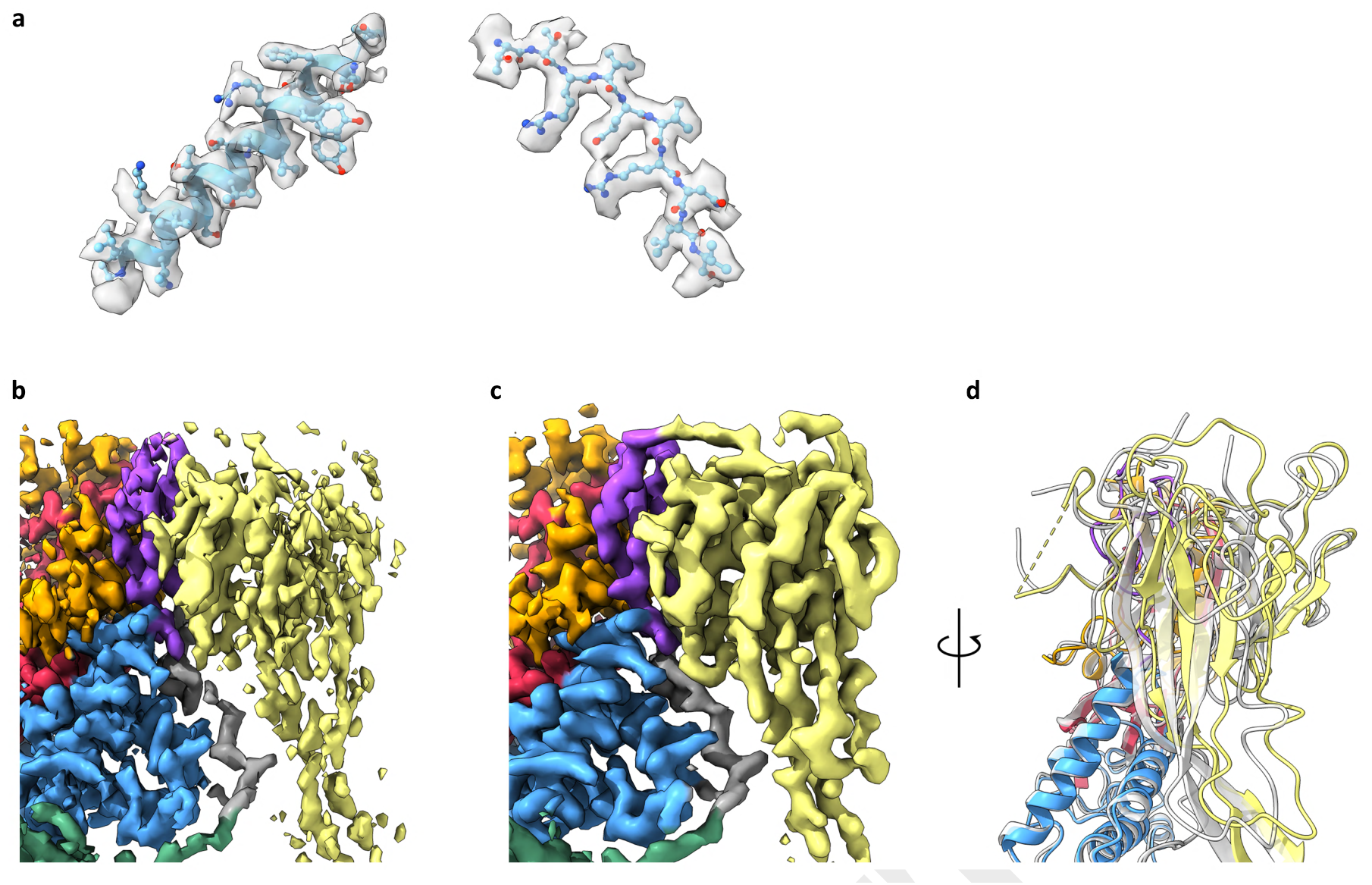
Conformational flexibility in the MABP domain. a) Select regions of electron density at the central core region of the MACPF domain from the consensus C16 reconstruction MPEG-1_*L*425*K*_ *α* showing high resolution features. b) The same reconstruction has poor, anisotropic resolution in the peripheral MABP domain, however after localised reconstruction c) the intramolecular heterogeneity can be resolved. d) Structural superposition of the two discrete conformations of the MABP domain.

**Fig. S8.**
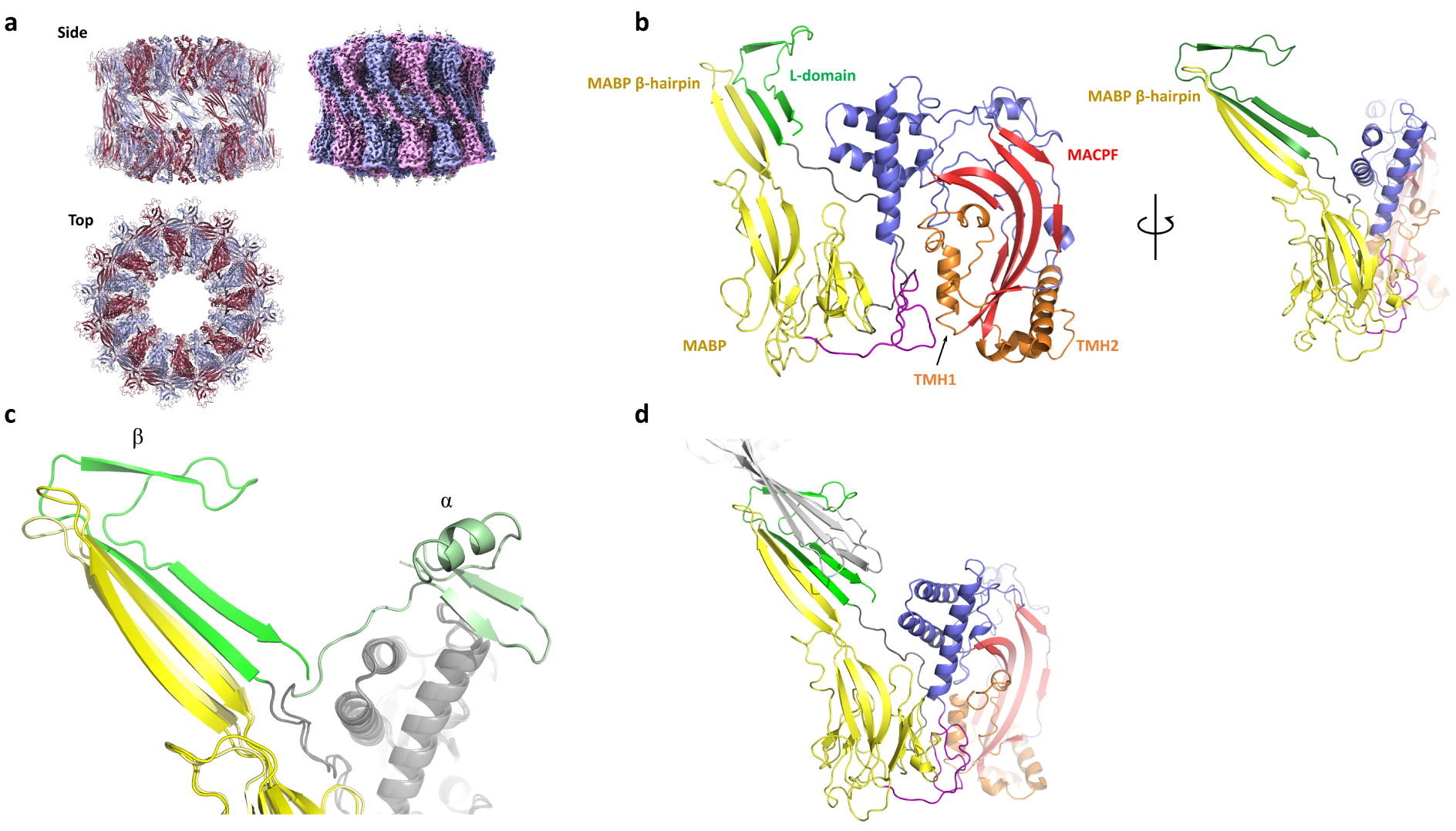
Structure (left) and cryoEM reconstruction (right) of the MPEG-1_*L*425*K*_ *β*-conformation assembly shown in alternating colours. b) The L-domain adopts a unique conformation resulting in a four stranded *β*-sheet extending from the original *β*-hairpin of the MABP domain. c) Rearrangement of the L-domain causes inter-subunit interactions to break resulting in d) *in trans* interactions between the two rings of the soluble head-to-head assembly.

**Table S1.**
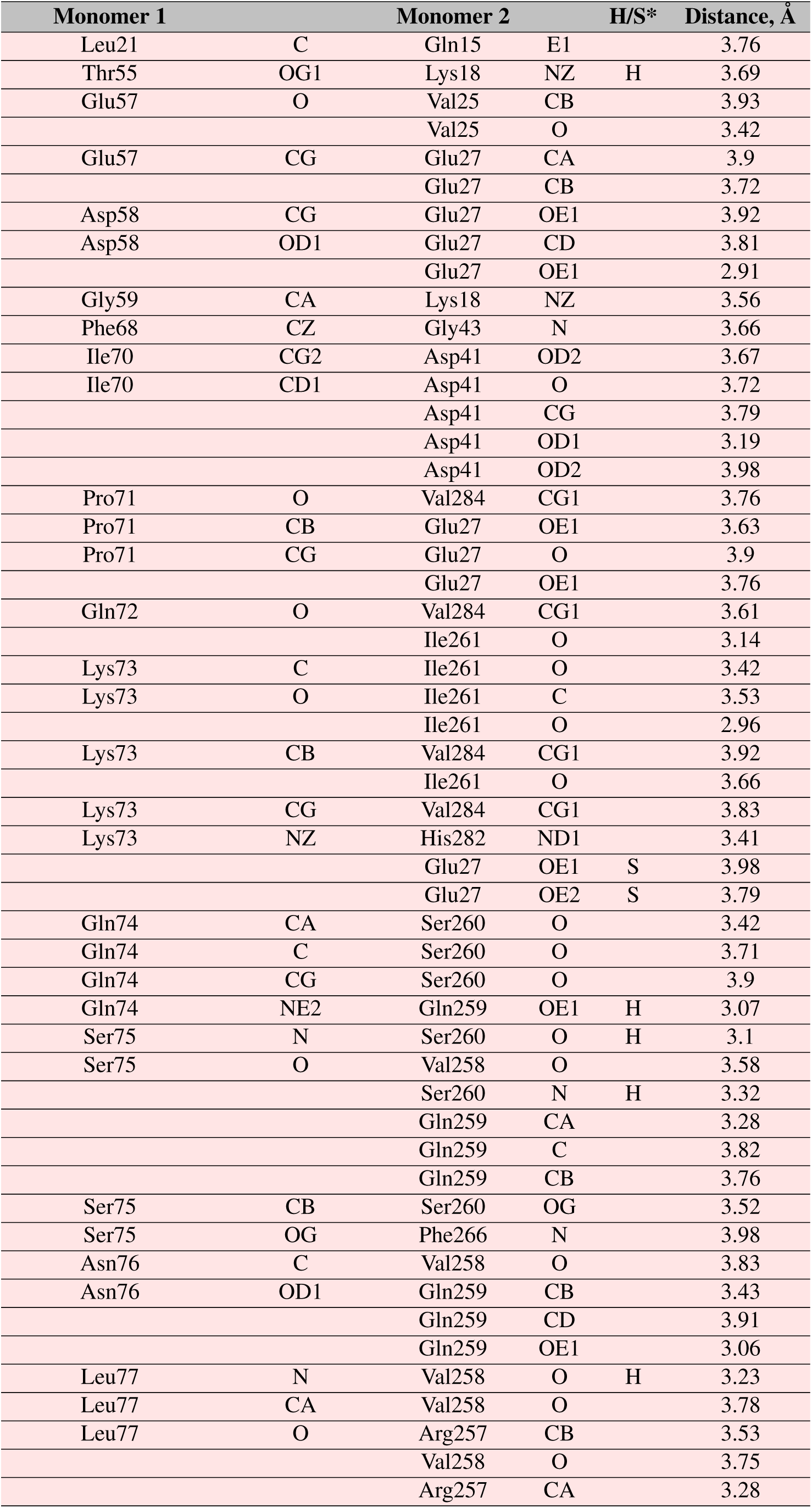
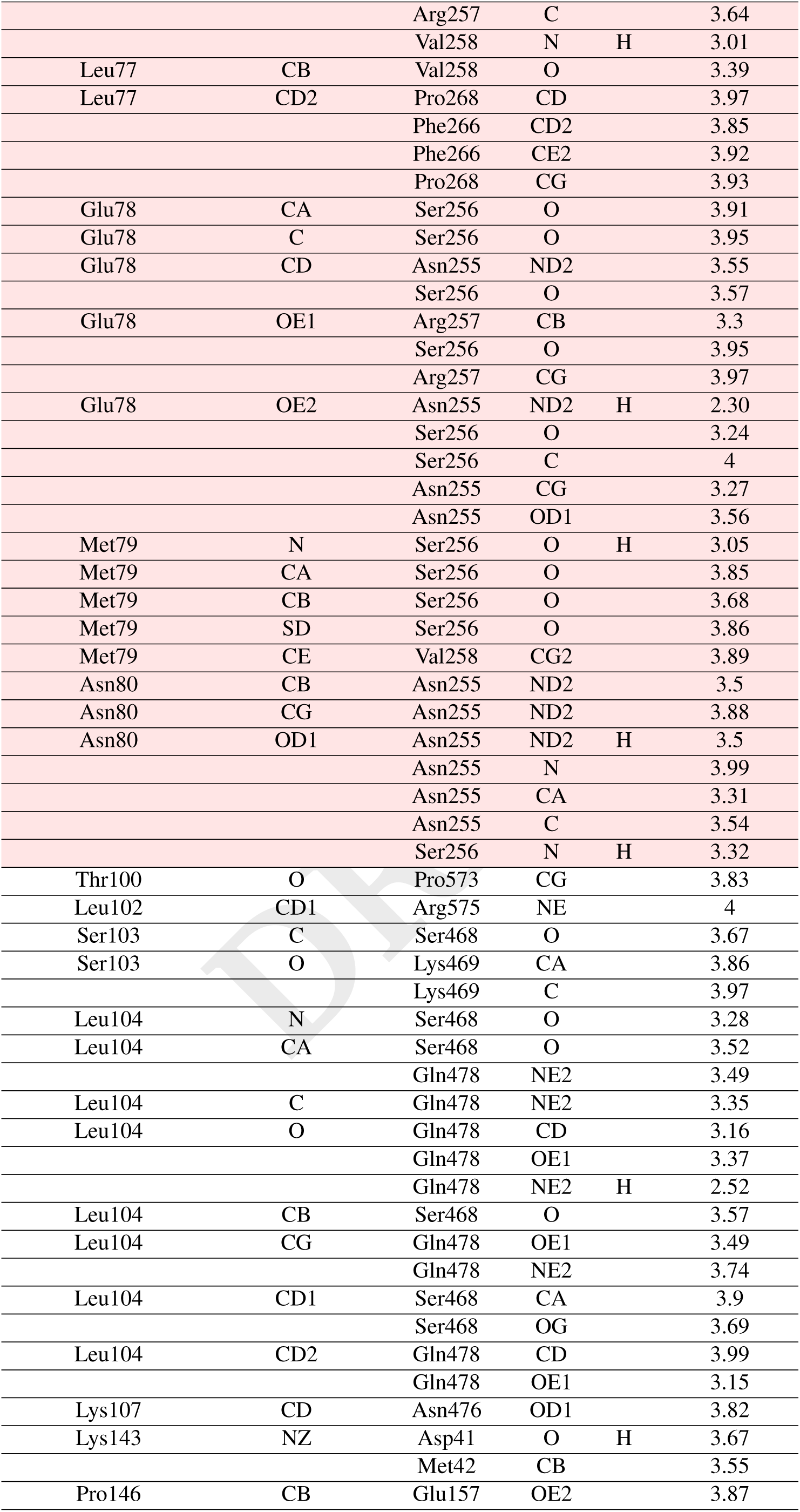
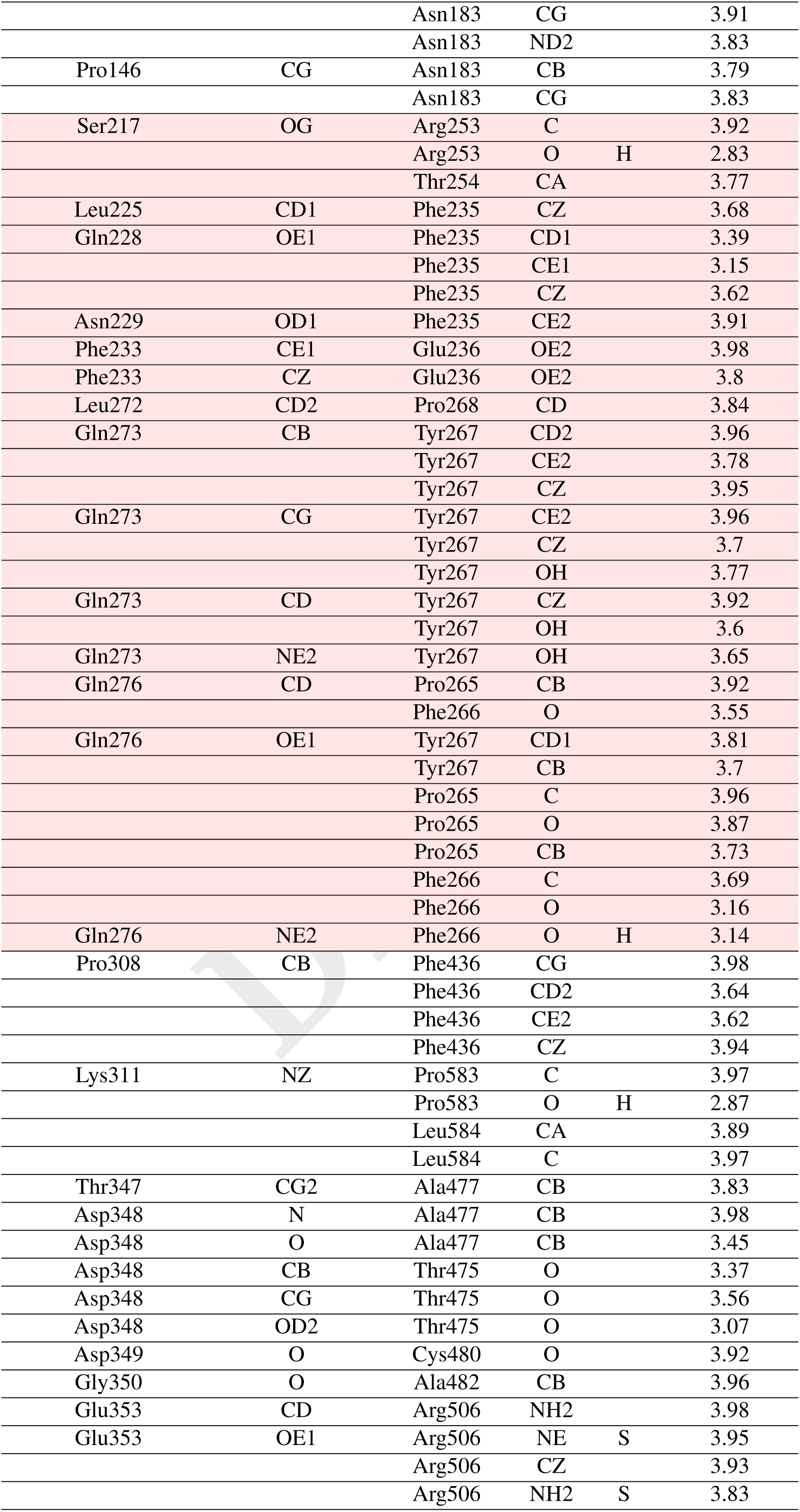
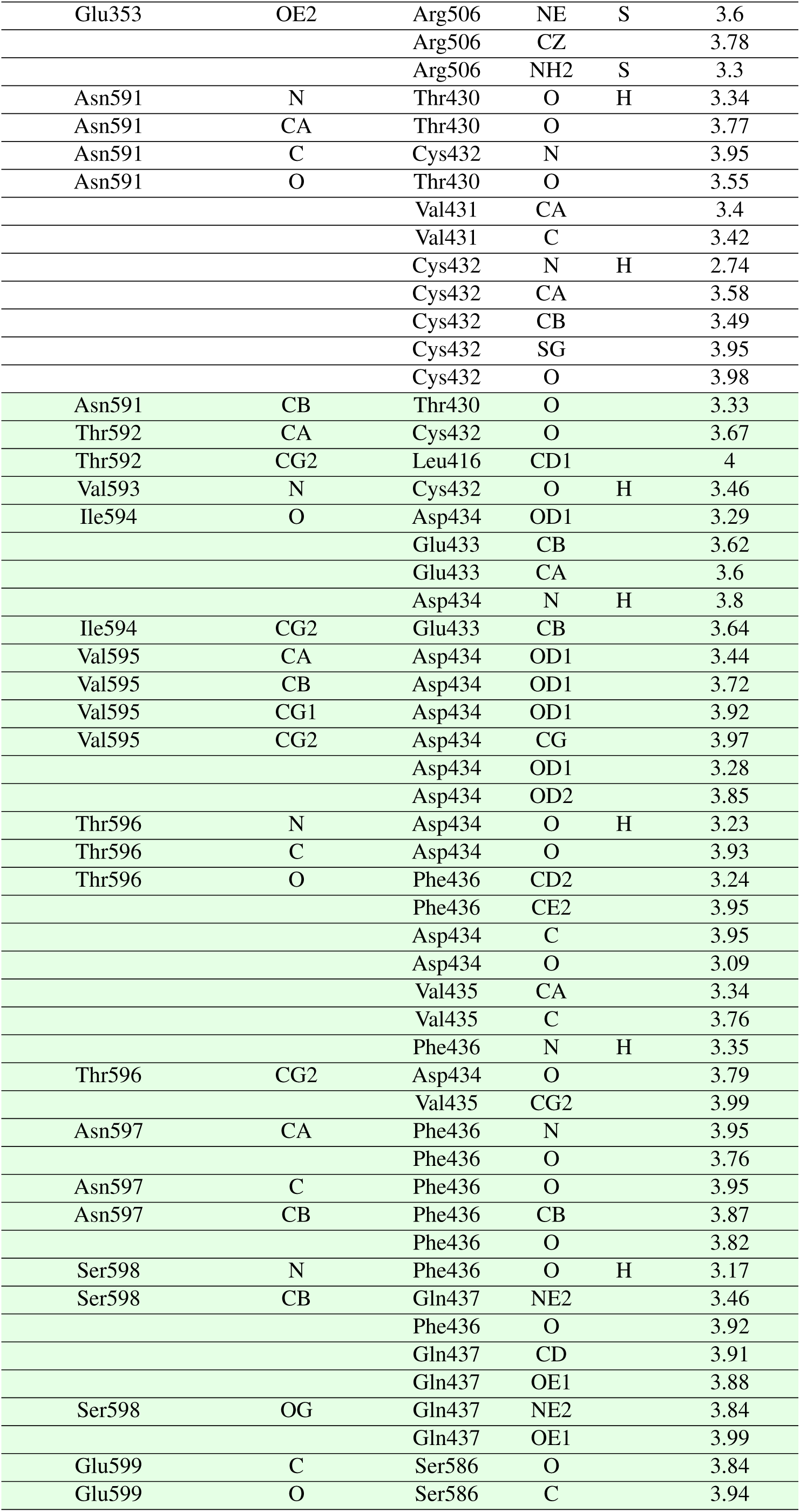
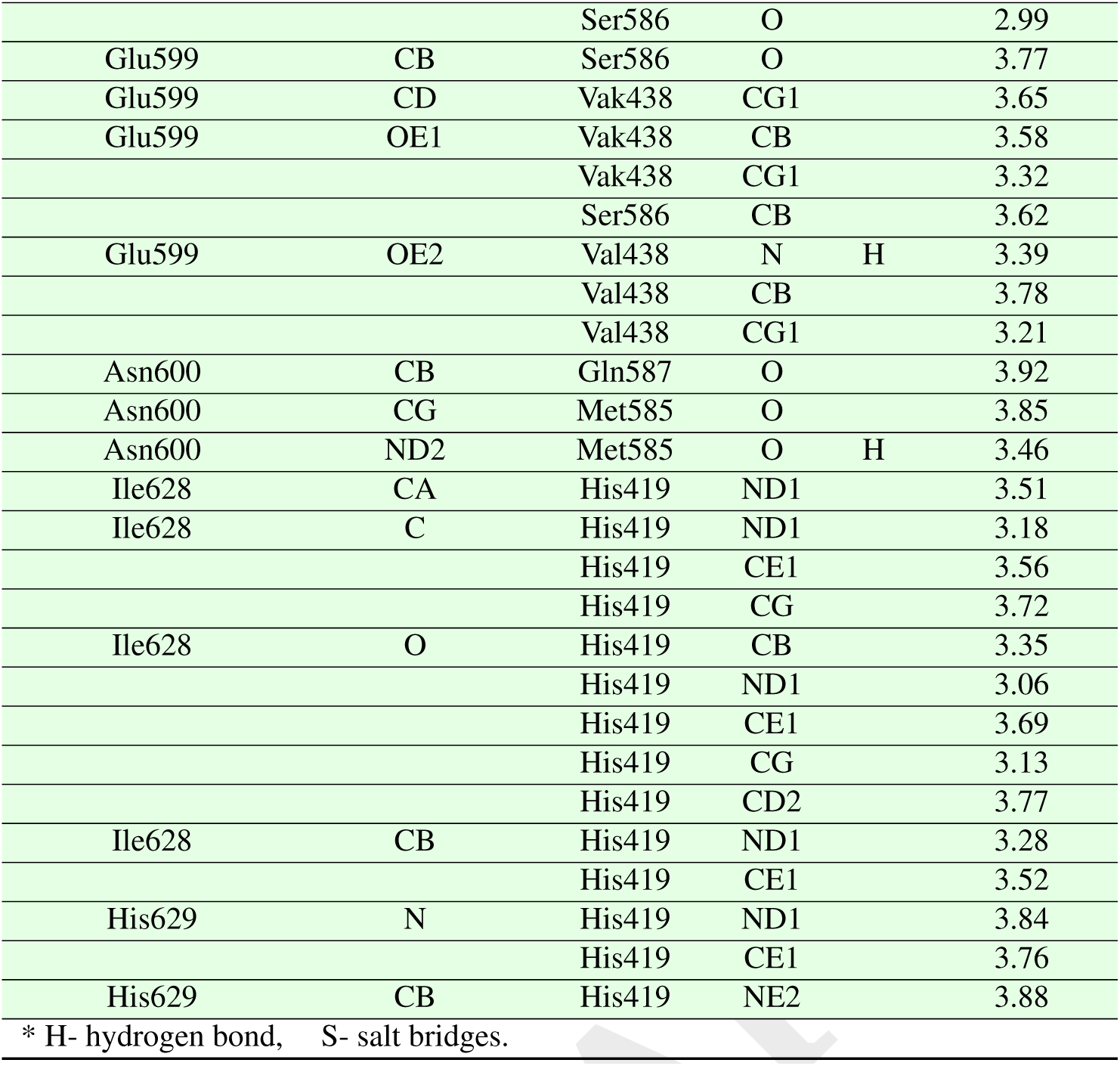
Contacts (≤ 4 Å) between adjacent MPEG-1 monomers (1 and 2) in the soluble MPEG-1 assembly (PDB XXX). The interface contacts were generated using CCP4(56)(CONTACT) and PISA(54). The contacts found around the central lumen of the *β*-barrel are shaded in red, and the *in trans* subunit interactions between the L-domain and the adjacent MABP *β*-hairpin are shaded in green.

**Table S2.**
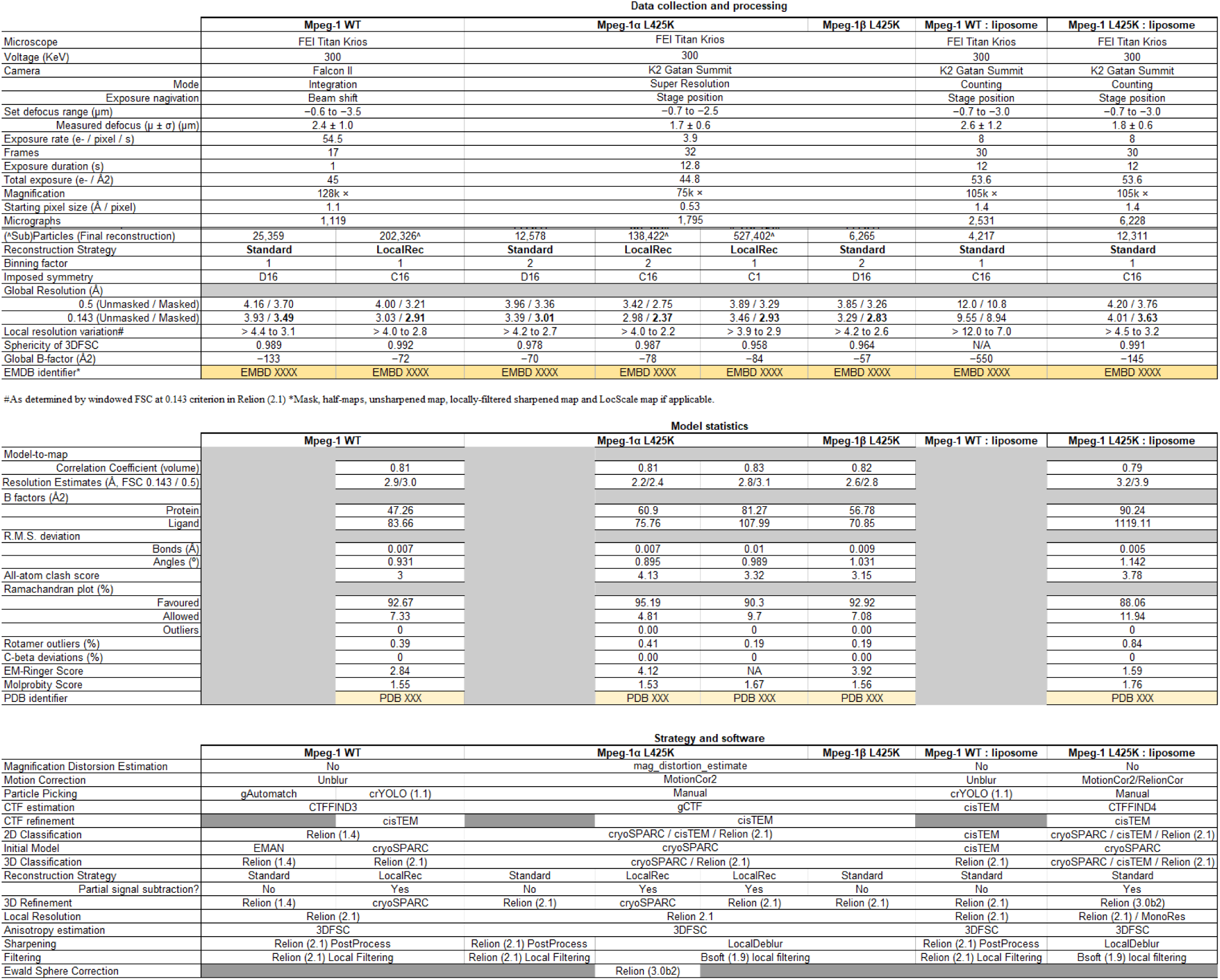
Summary table of Cryo-EM data collection, analysis and refinement statistics, including the particular software and processing strategy per data set.

